# New mathematical modelling tools for co-culture experiments: when do we need to explicitly account for signalling molecules?

**DOI:** 10.1101/2020.01.13.905414

**Authors:** Wang Jin, Haolu Wang, Xiaowen Liang, Michael S Roberts, Matthew J Simpson

## Abstract

Mathematical models are often applied to describe cell migration regulated by diffusible signalling molecules. A typical feature of these models is that the spatial and temporal distribution of the signalling molecule density is reported by solving a reaction–diffusion equation. However, the spatial and temporal distributions of such signalling molecules are not often reported or observed experimentally. This leads to a mismatch between the amount of experimental data available and the complexity of the mathematical model used to simulate the experiment. To address this mismatch, we develop a discrete model of cell migration that can be used to describe a new suite of co–culture cell migration assays involving two interacting subpopulations of cells. In this model, the migration of cells from one subpopulation is regulated by the presence of signalling molecules that are secreted by the other subpopulation of cells. The spatial and temporal distribution of the signalling molecules is governed by a discrete conservation statement that is related to a reaction–diffusion equation. We simplify the model by invoking a steady state assumption for the diffusible molecules, leading to a reduced discrete model allowing us to describe how one subpopulation of cells stimulates the migration of the other subpopulation of cells without explicitly dealing with the diffusible molecules. We provide additional mathematical insight into these two stochastic models by deriving continuum limit partial differential equation descriptions of both models. To understand the conditions under which the reduced model is a good approximation of the full model, we apply both models to mimic a set of novel co–culture assays and we systematically explore how well the reduced model approximates the full model as a function of the model parameters.

## 1 Introduction

Random motility is widely recognised as the key mechanism driving *in vitro* cell migration in highly idealised homogeneous environments (Huang et al. 2005; Treloar et al. 2014). However, in more realistic situations, cell migration is often regulated by external signals such as diffusible molecules. Cell migration regulated by signalling molecules plays an important role in embryonic development (Behar et al. 1996; Simpson et al. 2006), cancer metastasis (Kucia et al. 2004; Müller et al. 2001) and wound healing (Flegg et al. 2015; Pettet et al. 1996). In these situations, cell migration is often activated by signalling molecules binding to receptors on the cell surface (Yoon et al. 2016). Signalling molecules can be present in the environment or secreted by other cells (Luster 1998; Wright et al. 2005). In Figure 1(a) we show an example of such a system where a signalling molecule called stromal cell–derived factor 1 (SDF-1) binds to the C-X-C motif chemokine receptor 4 (CXCR4) expressed on the surface of a mesenchymal stem cell (MSC). This process can regulate migration of MSCs (Yoon et al. 2016).

**Fig. 1.**
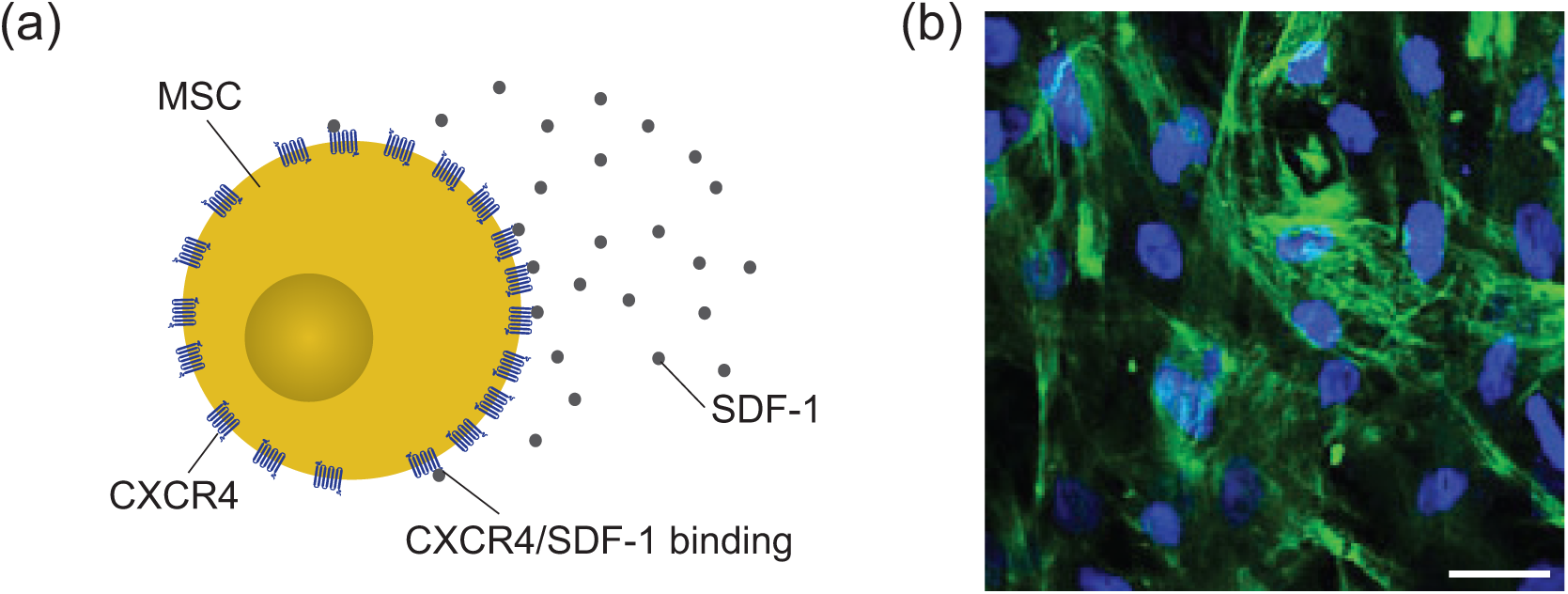
Binding of signalling molecules to biological cells. (a) A schematic showing CXCR4–SDF-1 binding on an MSC. (b) An immunofluorescence image of MSCs demonstrating the expression of CXCR4 which SDF–1 molecules bind to. The blue fluorescence indicates MSC nuclei. The green fluorescence indicates the expression of CXCR4. The scale bar corresponds to 50 *µ*m.

There are two key mechanisms that give rise to cell migration regulated by diffusible signalling molecules: (i) *chemokinesis* is where undirected cell migration is regulated by the local density of a particular signalling molecule (Liu and Klominek 2004; Cai et al. 2006); and (ii) *chemotaxis* is where the direction of cell migration is influenced by the spatial gradient of a signalling molecule (Keller and Segel 1971). The primary difference between chemokinesis and chemotaxis is that, at the individual level, chemokinesis influences the rate of undirected random cell movement without explicitly introducing a directional bias, whereas chemotaxis explicitly stimulates directional cell movement (Cai et al. 2006). Various experimental methods, such as transwell assays (Chen et al. 2006), microfluidic devices (Son et al. 2015), chemokinesis and chemotaxis assays (Richards et al. 2004; Rosoff et al. 2004), and co-culture migration assays (Chung et al. 2009; Frimberger et al. 2006) are used to study the role of chemokinesis and chemotaxis. However, these experimental approaches suffer from many important limitations. Two key limitations are: (i) signalling molecules are technically difficult to visualise in real time (Tokoyoda et al. 2004), and (ii) the spatial gradient of the signalling molecules is difficult to quantify (Chung et al. 2009).

Mathematical models have been widely used to mimic experimental observations relating to chemokinesis and chemotaxis (Brumley et al. 2019; Simpson et al. 2006). In the mathematical modelling literature, perhaps the most well–known model describing chemotaxis is the Keller–Segel partial differential equation (PDE) model proposed in 1971 (Keller and Segel, 1971). This fundamental continuum model has since been generalised to describe both chemokinesis and chemotaxis simultaneously (Balding and McElwain 1985; Byrne et al. 1998; Hillen and Painter 2009; Sherratt 1994). Further extensions of these continuum models include: (i) incorporating multiple cell populations (Stinner et al. 2014); (ii) explicitly modelling receptor–molecule binding (Sherratt et al. 1993); (iii) treating aggregates of cells as multiple interacting phases (Byrne and Owen 2004); and (iv) modelling responses with multiple signalling molecules (Painter et al. 2000). Apart from applying continuum PDEs to study cell migration stimulated by signalling molecules, discrete stochastic models have also been employed (Khain and Sander 2014; Pillay et al. 2018). Compared to continuum models, discrete models can be used to describe individual cell–level behaviour, and to specify how individual cells respond to signalling molecules. Using discrete models can be advantageous when comparing model predictions with experimental images that focus on individual cell level behaviour.

A key limitation of standard modelling frameworks is that typical models of chemokinesis and chemotaxis explicitly describe spatial and temporal distributions of the signalling molecules, often using a reaction–diffusion equation (Painter et al. 2000; Stinner et al. 2014). This is an important limitation because information about the spatial and temporal distributions of signalling molecules is rarely available from experiments (Chung et al. 2009; Tokoyoda et al. 2004). For example, we show an immunofluorescence image in Figure 1(b), with the green fluorescence indicating the expression of CXCR4. However, it is impossible to quantify the number of receptors or the spatial variability of the density of signalling molecules in this kind of standard experimental image. Therefore, it is unclear whether it is useful to mimic such an experiment with a mathematical model that explicitly describes the spatial and temporal variations in signalling molecule density. If one was to use a classical modelling approach, such as the Keller–Segel model, we would have no way of testing whether the spatial and temporal distributions of signalling molecules is accurate since these details are not available from standard experiments.

Motivated by new co–culture migration assays that we report in Section 2, the aim of this work is to develop an agent–based modelling framework that can be used to describe the dynamics of two interacting subpopulations of cells in a co–culture assay. In this model the movement of one type of agents is stimulated by the presence of signalling molecules that are produced by the other type of agents. The spatial and temporal distribution of the signalling molecules is governed by discrete conservation statement that is related to a reaction–diffusion PDE. We refer to this new model as the *full discrete model* since we explicitly describe the spatial and temporal distributions of agents and signalling molecules. To make the full discrete model more compatible with experimental data, we simplify the model by assuming that the dynamics of the signalling molecules is much faster than the time scale of cell migration. This simplification enables us to explore how the spatial distribution of signalling molecules affects cell movement without having to solve the underlying conservation equation for the signalling molecules. We refer to this simplified model as the *reduced discrete model*. The reduced model is both simpler to apply than the full discrete model since there are less parameters to estimate, as well as being more consistent with experimental observations in which the details of the signalling molecules are not reported. To provide additional mathematical insight into these two different stochastic models we also explore the continuum limit descriptions of both the full and reduced discrete models are derived through mean field analysis. This leads to new PDE models of signalling molecule–stimulated cell migration.

## 2 Co-culture experimental motivation

To motivate our modelling work we perform and report typical data from a series of *in vitro* co–culture ring barrier migration assays (Das et al. 2015). The full experimental protocol is documented in the Supplementary Material. Briefly, this type of co-culture assay involves uniformly seeding one type of cells inside a circular ring barrier and uniformly seeding another type of cells uniformly outside of the ring barrier (Das et al. 2015). Interactions between the two cell types can give rise to either a chemokinetic or chemotactic effect, depending upon the particular cell lines used in the experiment. In our expeiments, hepatocytes are seeded inside the ring barrier and MSCs are seeded outside the ring barrier in each experimental well in a 12–well tissue culture plate (Figure 2(a)–(b)). After seeding, the tissue culture plate is placed in an incubator overnight to allow cells to attach to the substrate. After attachment, the ring barrier is removed, leaving a vacant annulus of width approximately 1 mm (Figure 2(c)). Observations of the resulting cell migration are recorded by taking images of a small *field–of–view* over a 24 h period and recording the coordinates of particular cell trajectories over this period. Results in Figure 2(f)–(g) compare the endpoints of 20 typical MSC trajectories in two different experiments. The trajectories in Figure 2(f) are taken from a control experiment in which hepatocytes are omitted and we see that the MSCs appear to migrate randomly, with no obvious preferred direction. In contrast, the trajectories in Figure 2(g) are taken from an experiment that includes hepatocytes and we see clear evidence that the MSC migration is directed towards the location of the hepatocytes. Typical experimental results, such as those in Figure 2, do not provide any information about the temporal or spatial distribution of signalling molecules.

**Fig. 2.**
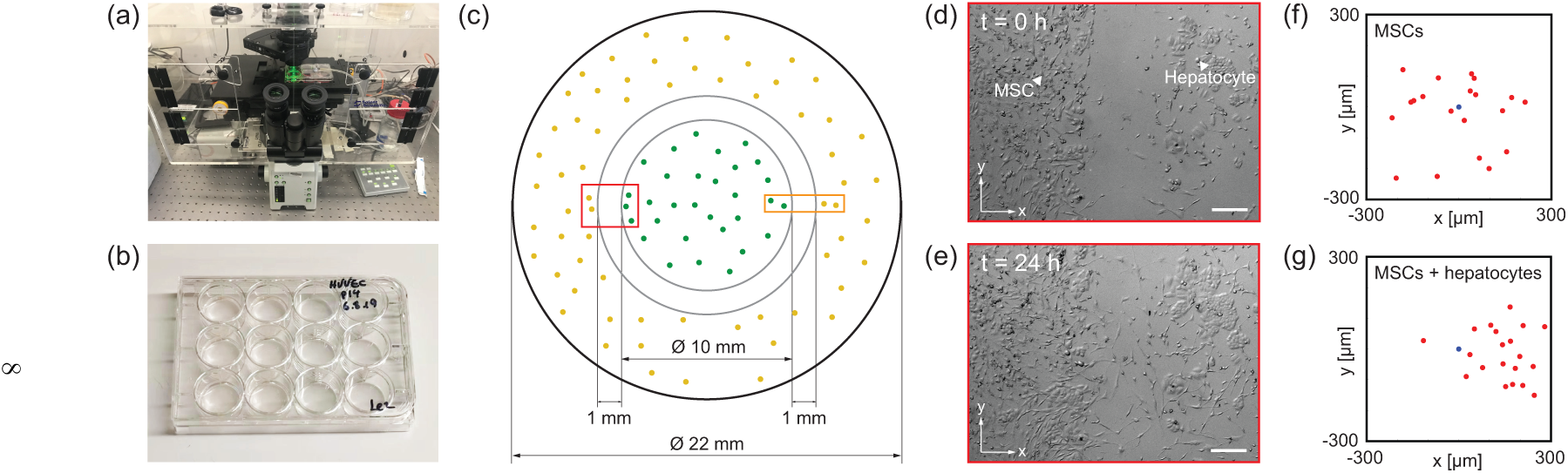
Ring barrier co-culture assays. (a) Live cell imaging microscope showing the incubator and confocal microscope apparatus. (b) An image of a 12-well plate. Each well has a diameter of 22 mm. (c) Schematic of a ring barrier migration assay. Initially hepatocytes (green dots) are placed uniformly inside the ring barrier and MSCs (yellow dots) are placed uniformly outside the ring barrier, leaving a vacant annulus highlighted in grey. The red rectangle indicates the field of view. The orange rectangle indicates the simulation domain.(d)–(e) Experimental image at *t* = 0 and 24h, respectively. The white scale bar corresponds to 200 *µ*m. (f) MSC trajectories in a control assay. (g) MSC trajectories in a co–culture assay including hepatocytes beyond the right boundary of the image. In both (f) and (g), the blue circles indicate cell positions at *t* = 0 h, and the red circles indicate cell positions at *t* = 24 h. All trajectories are shifted so that they start at the origin.

Many experimental studies indicate a role for chemokinesis or chemotaxis in co-culture experiments but do not show any spatial or temporal information about distributions of signalling molecules (Das et al. 2015). Therefore, we are motivated to model such experiments in a different way. For simplicity we apply our models to a small rectangular subregion as illustrated by the orange rectangle in Figure 2(c). Migration of cells in this subregion is predominantly horizontal, and to be consistent with this we develop our models in a one–dimensional geometry. Typical doubling times of MSC cell are over 50 h (Gruber et al. 2012), and since we only focus on relatively short time experiments we neglect the contribution of cell proliferation in our models.

## 3 Discrete models

The experimental data in Figure 2 provides strong evidence that MSC migration is biased in the presence of hepatocytes. Since hepatocytes are known to produce signalling molecules, such as SDF-1, we hypothesize that the directed migration of MCSs in Figure 2 is driven by a chemical signal. However, the data in Figure 2 does not indicate whether the directed migration arises from chemokinesis or chemotaxis, since both of these mechanisms can give rise to directed migration in the presence of a gradient of signalling molecule (Cai et al. 2006; Painter and Sherratt 2003). The modelling framework developed in this study can be used to examine either chemokinesis, chemotaxis, or a combination of chemokinesis and chemotaxis. For simplicity, we present the details by focusing on modelling chemokinesis in the main document. Additional results for modelling chemotaxis are presented in the Supplementary Material.

### 3.1 Full discrete model

We consider an agent–based model on a one–dimensional lattice where each site is indexed *i* ∈ [1, *I*] and has position *x* = (*i* − 1)Δ, where Δ is the lattice spacing that we take to be a typical cell diameter. The lattice is occupied by two different types of agents that represent the two different types of cells in the co-culture experiment: *Subpopulation 1* which secretes signalling molecules, such as the hepatocytes in Figure 2, and *Subpopulation 2* which senses and responds to the signalling molecules, such as the MSCs in Figure 2. The model is an exclusion process, meaning that each lattice site can be occupied by, at most, one agent. Therefore, in any single realisation of the model the occupancy of agents from Subpopulation 1 is given by *A*_*i*_ ∈ {0, 1}. If site *i* is occupied by an agent from Subpopulation 1 we have *A*_*i*_ = 1, and *A*_*i*_ = 0 otherwise. Similarly, in any single realisation of the model the occupancy of agents from Subpopulation 2 is given by *B*_*i*_ ∈ {0, 1}. The total number of agents from Subpopulation 1 and Subpopulation 2 are *N*_1_ and *N*_2_, respectively.

Since signalling molecules are many orders of magnitude smaller than cells, we allow each lattice site to be occupied by an arbitrary number of molecules, and we describe the density of signalling molecules at site *i* as *C*_*i*_ ∈ [0, ∞), where *C*_*i*_ is a continuous function of time. We assume that *C*_*i*_ is measured in some appropriate unit, such as *µ*M. Such hybrid models that treat cells as discrete objects and signalling molecules as continuous densities is standard in the mathematical biology literature (Alacón et al. 2003; Mallet and de Pillis 2006). We will now specify how agents on the lattice move in response to the signalling molecules.

#### 3.1.1 Agent movement

Within a particular time step of duration *τ*, agents from Subpopulation 1 and Subpopulation 2 attempt to undergo a nearest neighbour random walk with probability *f*_1_(*C*_*i*_) ∈ [0, 1] and *f*_2_(*C*_*i*_) ∈ [0, 1], respectively. The functional forms of *f*_1_(*C*_*i*_) and *f*_2_(*C*_*i*_) determine how the agents respond to the density of signalling molecules. For example, if *f*_1_(*C*_*i*_) is increasing, the signalling molecules amplify the migration rate of Subpopulation 1. We will discuss the particular choice of *f*_1_(*C*_*i*_) and *f*_2_(*C*_*i*_) in Section 5.2.

Agent movement is simulated using a random sequential update method. During a time step of duration *τ, N*_1_ + *N*_2_ agents are randomly selected, one at a time, with replacement (Jin et al. 2016a). If an agent from Subpopulation 1 at site *i* is selected, that agent attempts to undergo a nearest neighbour random walk with probability *f*_1_(*C*_*i*_). Similarly, if an agent from Subpopulation 2 at site *i* is selected, that agent attempts to undergo a nearest neighbour random walk with probability *f*_2_(*C*_*i*_). In all cases the target site is chosen at random, and potential motility events are aborted if the target site is occupied. Reflecting boundary conditions are applied.

#### 3.1.2 Signalling molecules

To be consistent with our experimental observations in Figure 1, we assume the signalling molecules are secreted by agents from Subpopulation 1 at a particular rate. The signalling molecules diffuse, undergo decay, and are taken up by agents from Subpopulation 2. We suppose that the spatial and temporal distribution of signalling molecules is governed by discrete conservation statement,

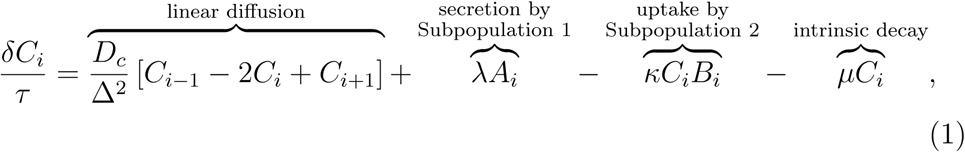

where *D*_*c*_ [*µ*m^2^/h] is the molecular diffusivity, *λ* [*µ*M/h] is the secretion rate, *κ* [/h] is the uptake rate and *µ* [/h] is the intrinsic decay rate. We solve Equation (1) numerically as outlined in the Supplementary Material.

### 3.2 Reduced discrete model

We now formulate a reduced discrete model that retains key elements of the full discrete model without the need to explicitly solve for the spatial and temporal distribution of the signalling molecules. To distinguish between the two models we write the site occupancy of Subpopulation 1 and Subpopulation 2 as *U*_*i*_ ∈ {0, 1} and *V*_*i*_ ∈ {0, 1}, respectively. The diffusivity of signalling molecules is approximately three orders of magnitude greater than a typical cell diffusivity (Jin et al. 2017; Mac Gabhann and Popel 2004). This motivates us to simplify the model by assuming we have quasi-steady conditions since the diffusive transport evolves on much faster timescale than the source terms on the right of Equation (1). If the magnitude of the source terms in Equation (1) are negligible relative to the diffusive transport term, at steady state we have *C*_*i*+1_ − 2*C*_*i*_ + *C*_*i*−1_ = *δC*_*i*_ = 0. Setting *C*_*i*+1_ − 2*C*_*i*_ + *C*_*i*−1_ = 0 and *δC*_*i*_ = 0 in Equation (1) gives *C*_*i*_ = *λU*_*i*_*/*(*µ* + *κV*_*i*_), which could be a useful way to indirectly represent the effect of the signalling molecules as a function of the spatial arrangement of the agents on the lattice. This kind of quasi-steady assumption is often used to simplify continuum mathematical models where some kind of diffusible signal (e.g. Cai et al. 2006) or diffusible nutrient (e.g. Breward et al. 2002) is assumed to approach steady state much faster than the dynamics of some population of cells. The consequences of making such assumptions in a stochastic framework are rarely, if ever, examined in detail.

Since we have an exclusion process, each lattice site can be occupied by a single agent. Therefore, simply applying *C*_*i*_ = *λU*_*i*_*/*(*µ* + *κV*_*i*_) leads to *C*_*i*_ = 0 at any site with *U*_*i*_ = 0, or *C*_*i*_ = *λ/µ* for any site with *U*_*i*_ = 1. To make this approximation more realistic, we take the occupancy of lattice site *i* to be the average of the nearest neighbour lattice sites, *Û*_*i*_ = (*U*_*i*−1_ + *U*_*i*+1_)/2 and 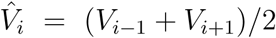, giving

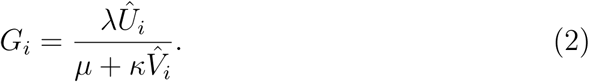

Therefore, in the reduced model, we take *G*_*i*_ to approximate density of the signalling molecule density at site *i*. Using this approximation in our discrete modelling framework allows us to implicitly simulate the role of the signalling molecules without needing to solve Equation (1). This approach has three clear advantages over the full discrete model: (i) the reduced discrete model involves less parameters than the full discrete model; (ii) the reduced discrete model is faster to computer than the full discrete model since there is no need to solve the evolution equation for *C*_*i*_; and (iii) the reduced discrete model is more consistent with typical experimental observations that do not measure or report spatial and temporal distributions of the signalling molecules.

The reduced discrete model is implemented computationally using a similar random sequential update method. The only difference is that in the reduced discrete model we apply *f*_1_(*G*_*i*_) and *f*_2_(*G*_*i*_) instead of *f*_1_(*C*_*i*_) and *f*_2_(*C*_*i*_), and there is no need to solve the evolution equation for *C*_*i*_. Of course, the key question that we are interested in now is to establish when the reduced model provides a good approximation to the full model. Intuitively we expect that the reduced discrete model will be a good approximation of the full model when *C*_*i*_ is accurately approximated by *G*_*i*_. However, to explore this quantitatively we need to compare the performance of the two models over a series of biologically relevant parameter values. Before we consider this comparison, we also provide more mathematical insight into the two models by deriving approximate continuum limit descriptions of the two discrete modelling frame-works.

## 4 Continuum limit descriptions

We begin the continuum limit derivation by assuming we have access to a large number of identically prepared realisations of the full discrete model, and we denote the average occupancy of Subpopulation 1 and Subpopulation 2 at site *i* by *Ā*_*i*_ ∈ [0, 1] and 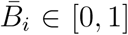, respectively. Similarly, the average density of signalling molecules at site *i* is given by 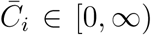). Invoking a mean-field assumption and accounting for all possible events that alter the occupancy of site *i* over a time step of duration *τ*, we obtain

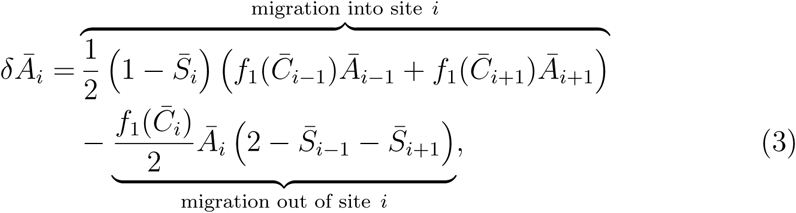

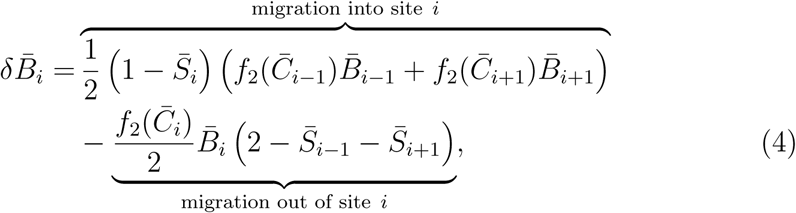

where *δĀ*_*i*_ and 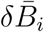 are the change in occupancy at site *i* of Subpopulation 1 and 2, respectively, and 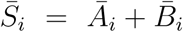 is the total average occupancy at site *i*. To convert these discrete conservation statements into continuous expressions we identify the discrete variables with appropriate continuous variables, *Āi*(*t*) = *a*(*x, t*), 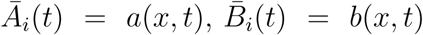 and 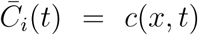. Expanding each term in Equations (3)–(4) about site *i* using a Taylor series and neglecting terms of *𝒪*(Δ^3^), we divide both sides of the resulting expressions by *τ* and take the limit Δ → 0 and *τ* → 0 jointly, with the ratio Δ^2^*/τ* held constant, to give

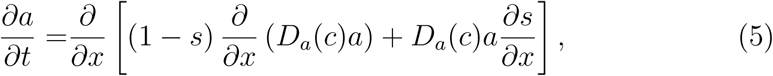

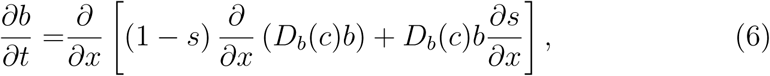

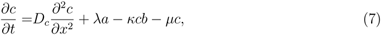

where *D*_*a*_ = Δ^2^*f*_1_(*c*)*/* (2*τ*) and *D*_*b*_ = Δ^2^*f*_2_(*c*)*/* (2*τ*) are the diffusion coefficients for Subpopulation 1 and Subpopulation 2, respectively and *s*(*x, t*) = *a*(*x, t*)+ *b*(*x, t*). We refer to Equations (5)–(7) as the *full continuum model*.

The continuum limit description of the reduced discrete model can be obtained using a very similar approach. The approximate conservation statements for the two subpopulations can be written as,

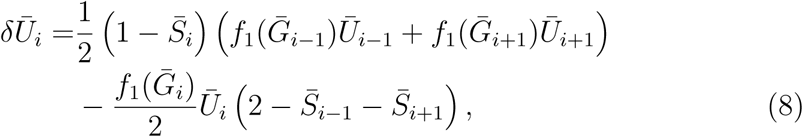

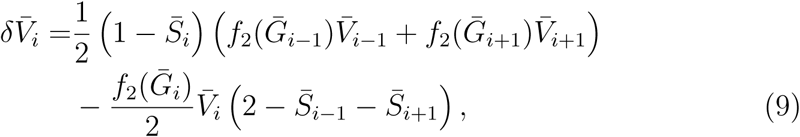

where all terms have a similar interpretation to those in Equations (3)–(4). We proceed to the continuum limit in the same way, arriving at

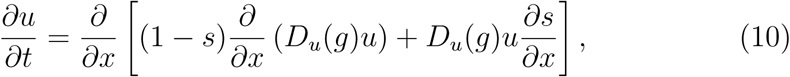

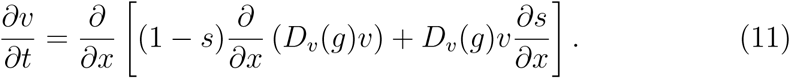

where *u*(*x, t*) and *v*(*x, t*) are the densities of Subpopulation 1 and Subpopulation 2, respectively. Here, *D*_*a*_ = Δ^2^*f*_1_(*g*)*/* (2*τ*) and *D*_*b*_ = Δ^2^*f*_2_(*g*)*/* (2*τ*) are the diffusion coefficients for Subpopulation 1 and Subpopulation 2, respectively. We refer to Equations (10)–(11) as the *reduced continuum model*.

## 5 Results and Discussion

In this section we explore solutions of the full and reduced models, both discrete and continuum, for a typical geometry and timescale that reflect the co–culture assay in Figure 2. For the discrete models we set *τ* = 0.01 h and Δ = 20 *µ*m to reflect a typical cell diameter. To simulate the width of the experimental field-of-view in Figure 2(c) we choose *I* = 201. We initialise the discrete simulations by placing agents from Subpopulation 1 to the left of the domain and agents from Subpopulation 2 to the right of the domain at *t* = 0. All sites with *i ≤* 76 are randomly populated with probability 0.6 by agents from Subpopulation 1 and all sites with *i* ≥ 126 are randomly populated with probability 0.6 by agents from Subpopulation 2. This initial condition leaves 1000 *µ*m of vacant space in the middle of the domain which is consistent with the initial width of the annulus of free space in Figure 2. In the full discrete model we assume that *C*_*i*_ = 0 at all sites at *t* = 0.

The full and reduced continuum models are solved numerically as outlined in the Supplementary Material. The initial condition in the continuum model is consistent with the discrete models by setting *a*(*x*, 0) = 0.6 for 0 *≤ x ≤* 1500 *µ*m, *b*(*x*, 0) = 0.6 for 2500 *≤ x ≤* 4000 *µ*m, and *a*(*x*, 0) = *b*(*x*, 0) = 0 elsewhere. In the full continuum model we set *c*(*x*, 0) = 0 for 0 *≤ x ≤* 4000 *µ*m and we will comment on this choice of initial conditions later.

### 5.1 Choice of model parameters

To make our simulations consistent with experimental observations we note that MSCs are known to respond to diffusible molecules secreted by hepatocytes in co–culture assays, whereas the migration of hepatocytes are unaffected by the presence of MSCs in co–culture (Novo et al. 2011; Yoon et al. 2016). Accordingly, in the discrete models we assume that agents from both subpopulations undergo unbiased migration when *C*_*i*_ = 0 and that *C*_*i*_ has no impact upon the migration of Subpopulation 1 so we set *f*_1_(*C*_*i*_) to be a constant. In contrast, we choose *f*_2_(*C*_*i*_) to be a smooth increasing function, given by

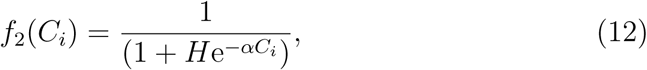

where *α* ≥ 0 specifies the strength of the chemokinetic response, and *H* is a constant relating to the migration rate of Subpopulation 2 in the absence of signalling molecules. In this section we choose *H* = 9, which gives *f*_2_(0) = 1*/*10. We set *f*_1_(*C*_*i*_) = *f*_2_(0) = 1*/*10 so that in the absence of the chemical signal, agents from both subpopulations undergo unbiased random migration at the same rate. In terms of the continuum limit description, our choices of Δ, *τ, f*_1_(*C*_*i*_) and *f*_2_(*C*_*i*_) correspond to *D*_*a*_ = *D*_*b*_(0) = *D*_*u*_ = *D*_*v*_(0) = 2000 *µ*m^2^/h which is a typical value of cell diffusivity in low density tissue culture (Jin et al. 2016b).

There are five free parameters in the full and reduced models: *D*_*c*_, *λ, κ, µ*, and *α*. We note that the diffusivity of typical diffusible molecules is approximately 10^5^ *µ*m^2^/h (Mac Gabhann and Popel 2004; Veldkamp et al. 2009). Experimental observations of the half life of diffusible molecules is around 0.5 h (Kirkpatrick et al. 2010), which corresponds to an exponential decay rate of approximately 1 /h. Therefore, we set *D*_*c*_ = 10^5^ *µ*m^2^/h and *µ* = 1 /h. We are unaware of any detailed experimental measurements of production and uptake rates of SDF-1 for co–culture experiments with hepatocytes and MSC so we choose *λ* = 1 *µ*M/h and *κ* = 1 /h, to be of the same order as the decay rate. Later we will vary these choices of parameter values to gain insight into the sensitivity of the model predictions to these choices of parameter values.

### 5.2 Comparisons of the full and reduced models

Results in Figure 4(a)–(b) show snapshots of the time evolution of agent positions in the full and reduced discrete models, respectively. In these preliminary simulations we specify a weak chemokinetic effect, *α* = 1. Comparing the distribution of agents in different rows of the subfigures shows that the two subpopulations migrate into the initially–vacant space over time. We estimate the expected behaviour of the simulations by averaging the occupancy of each lattice site using 500 identically–prepared realisations of the stochastic models and show the averaged density profiles in Figure 4(c) where we see that the averaged density profiles from the reduced discrete model compares very well.

**Fig. 3.**
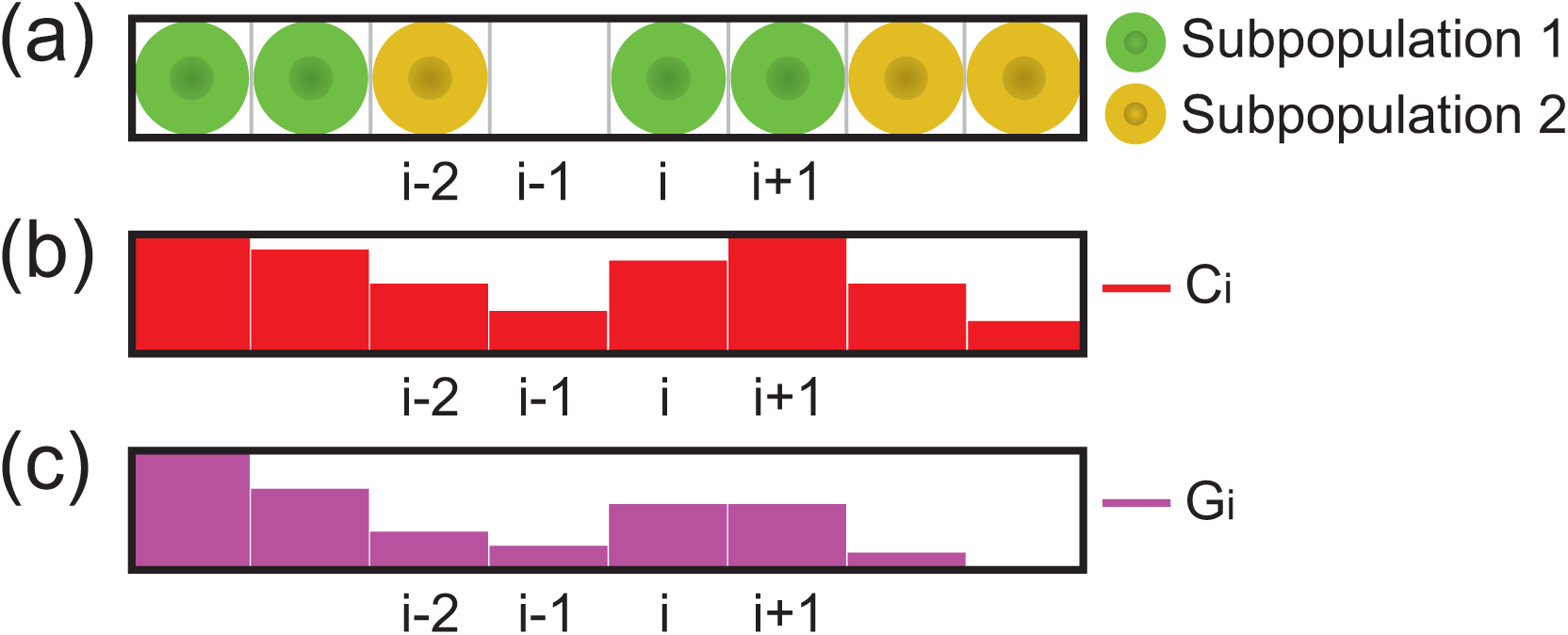
Discrete modelling framework. (a) A schematic of the agent–based model comprising Subpopulation 1 and Subpopulation 2. (b) Spatial distribution of the signalling molecule density in the full discrete model. (c) Spatial distribution of the approximate density of the signalling molecule density in the reduced discrete model.

**Fig. 4.**
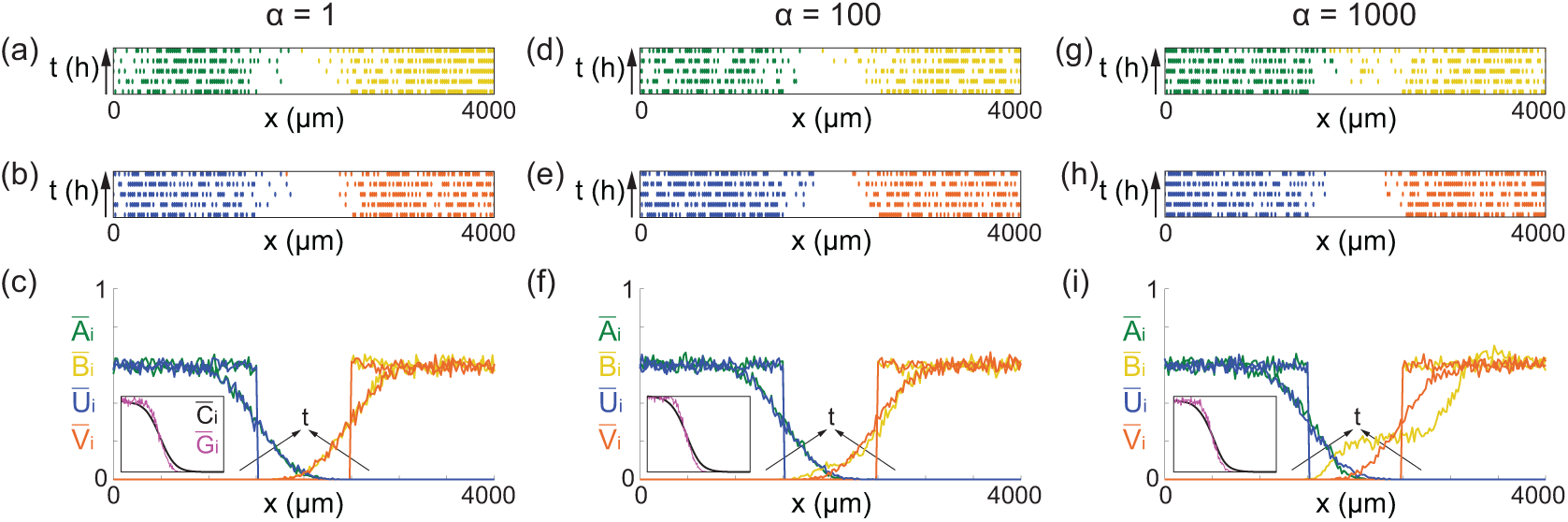
Comparison of the full and reduced discrete models. (a), (d), (g) Snapshots from the full discrete model at *t* = 0, 6, 12, 18, and 24 h. The arrow along the vertical direction indicates increasing time. (b), (e), (h) Snapshots from the reduced discrete model at *t* = 0, 6, 12, 18, and 24 h. (c), (f), (i) Density profiles of Subpopulation 1 and Subpopulation 2 from the full and reduced discrete models at *t* = 0 and 24 h. The black arrow indicates increasing time. The inset in each subfigure shows profiles of the 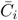 and 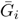 at *t* = 24 h. All the simulation data are obtained by averaging over 500 statistically identically prepared realisations. *D*_*a*_ = *D*_*b*_(0) = *D*_*u*_ = *D*_*v*_(0) = 2000 *µ*m^2^/h, *D*_*c*_ = 10^5^ *µ*m^2^/h, *λ* = 1 *µ*M/h, and *κ* = *µ* = 1 /h, Δ = 20 *µ*m and *τ* = 0.01 h for all the simulations.

To investigate how the comparison between the full and reduced discrete models depends upon the strength of the chemokinetic effect we present additional results in the in Figure 4 (d)–(e) for *α* = 100 and in Figure 4 (g)–(h) for *α* = 1000. Comparing the averaged density profiles at *t* = 24 h shows that we maintain reasonably good agreement between the reduced and full models for the moderate chemokinetic effect in Figure 4(f) but we see that the reduced discrete model does not approximate the full discrete model very well when the chemokinetic effect is strong, as in Figure 4(i). In addition, in the Supplementary Material we compare the averaged density profiles at *t* = 48 h to allow more time, beyond the typical experimental timescale, for the two subpopulations to interact. These additional results over a longer time scale are consistent with the results in Figure 4.

All results in Figure 4 correspond to discrete results. We now examine how well the averaged data from the two discrete models compare with the numerical solution of the associated continuum limit descriptions. Results in Figure 5(a)–(c) compare averaged density profiles from the full model with corresponding solutions of Equations (5)–(7) for *α* = 1, 100 and 1000, respectively. These results show that the new PDE models provide an accurate approximation of the averaged behaviour of the full discrete model when *α* = 1 and *α* = 100, but that the solution of the continuum limit PDE does not provide an accurate approximation of the averaged data from the full discrete model when chemokinesis is sufficiently strong, *α* = 1000. Similarly, results in Figure 5(d)–(f) compare averaged density profiles from the reduced model with corresponding solutions of Equations (10)–(11) for *α* = 1, 100 and 1000, respectively. Again, we see that the solution of the continuum limit PDE models provides a good approximation of the averaged behaviour of the reduced discrete model when *α* = 1 and *α* = 100, but we observe some discrepancy when the chemokinesis is sufficiently strong, *α* = 1000. Therefore, while the continuum limit PDEs can provide a good description of the average behaviour of the discrete model for certain parameter choices, they do not always provide a good approximation of the discrete models and this discrepancy is associated with the failure of the mean-field approximation (Simpson et al. 2010). Therefore, for the remainder of this study we will focus on using the discrete models and explore the differences in the performance of the full and reduced discrete models.

**Fig. 5.**
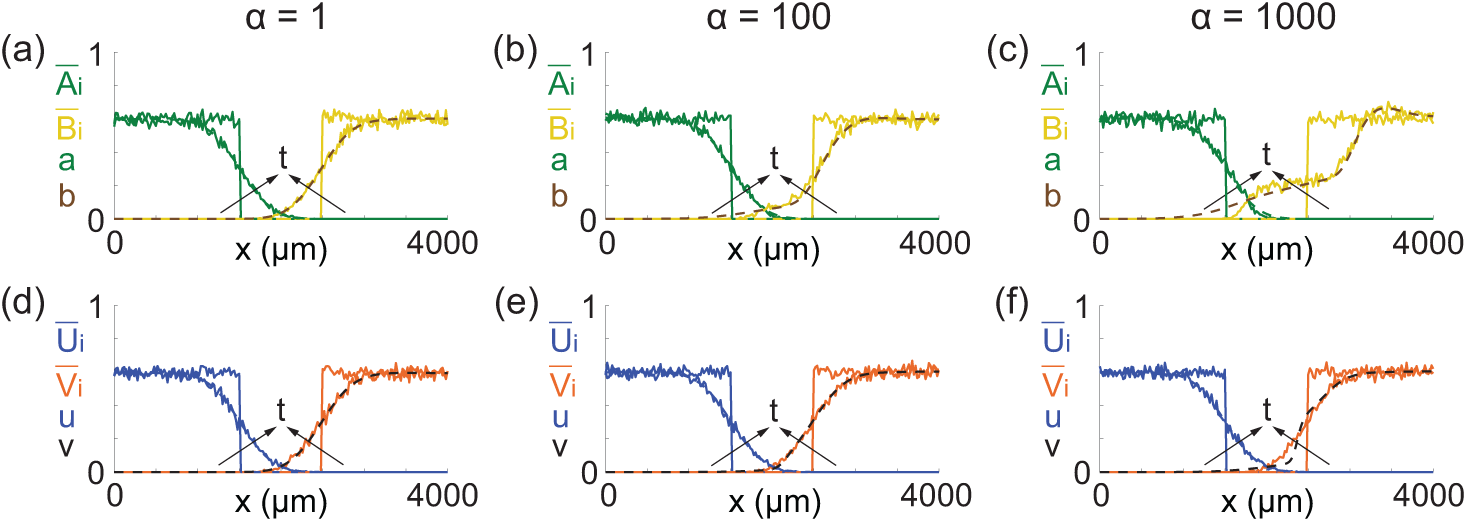
Continuum–discrete comparisons of the full and reduced models. (a)–(c) Continuum–discrete comparisons of the full models at *t* = 0 and 24 h. (d)–(f) Continuum–discrete comparisons of the reduced models at *t* = 0 and 24 h. The solid line indicates results from the discrete models. The dashed line indicates results from the continuum models. The black arrow indicates increasing time. *D*_*a*_ = *D*_*b*_(0) = *D*_*u*_ = *D*_*v*_(0) = 2000 *µ*m^2^/h, *D*_*c*_ = 10^5^ *µ*m^2^/h, *λ* = 1 *µ*M/h, and *κ* = *µ* = 1 /h for all the cases. Δ = 20 *µ*m and *τ* = 0.01 h for all the discrete simulations.

We now quantitatively explore the difference between the full and reduced discrete models for a range of signalling molecules diffusivity (*D*_*c*_ = 10, 10^5^, 10^6^ *µ*m^2^/h) and a range of chemokinetic strengths (*α* = 1, 100, 1000). To quantify the quality-of-match between the full and reduced models, we compute a measure of the least–squares difference between the averaged density profiles,

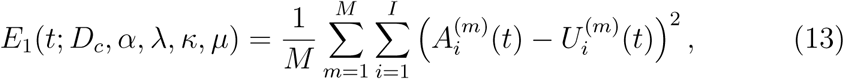

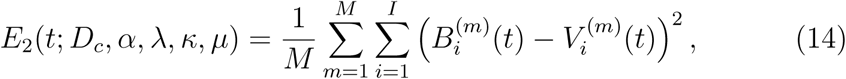

where *m* is an index indicating the number of identically-prepared realisations and *M* = 500 is the total number of identically–prepared realisations considered. For each combination of *D*_*c*_ and *α* that we consider, we compute *E*_1_(24; *D*_*c*_, *α, λ, κ, µ*) and *E*_2_(24; *D*_*c*_, *α, λ, κ, µ*) with fixed values of *λ* = 1 *µ*M/h and *κ* = *µ* = 1 /h, and we plot the averaged density profiles at *t* = 24 h in Figure 6. Results in Figure 6 indicate that *E*_1_ is relatively small and insensitive to the parameter values we consider. In contrast, *E*_2_ increases with both *α* and *D*_*c*_. In particular we see that the reduced discrete model can provide a very good approximation of the full discrete model when *α* is sufficiently small, but the approximation becomes poor when *α* increases.

**Fig. 6.**
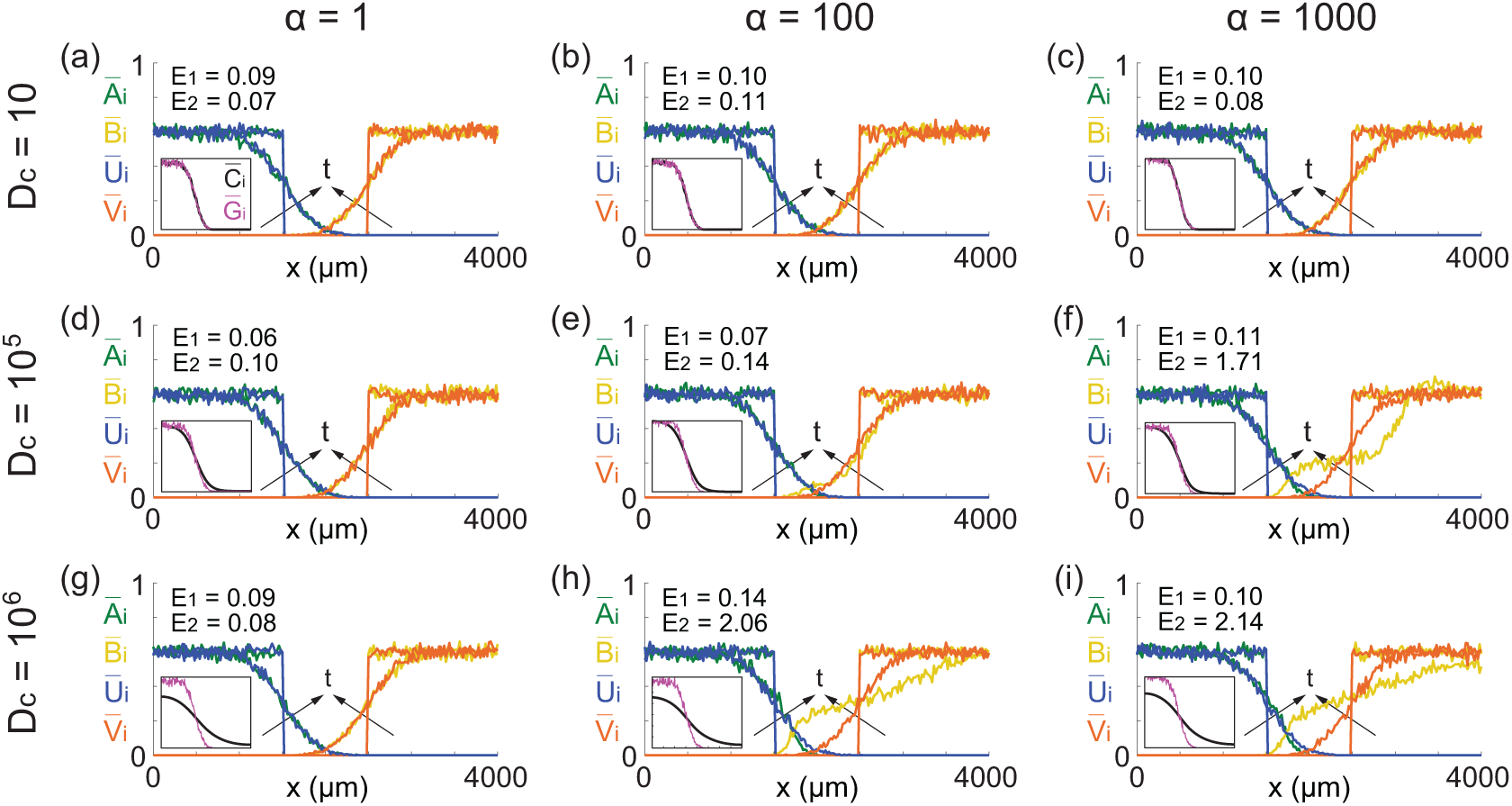
Comparisons of the results form the full and reduced discrete models. Density profiles of Subpopulation 1 and Subpopulation 2 from the full and reduced discrete models at *t* = 0 and 24 h are shown. The inset in each subfigure compares 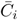 and 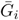 at *t* = 24 h. The black arrow indicates increasing time. *D*_*a*_ = *D*_*b*_(0) = *D*_*u*_ = *D*_*v*_(0) = 2000 *µ*m^2^/h, *λ* = 1 *µ*M/h, *κ* = *µ* = 1 /h, Δ = 20 *µ*m and *τ* = 0.01 h for all the simulations.

In addition to comparing averaged agent density profiles for the full and discrete models, the insets provided in each subfigure of Figure 6 show the spatial distributions of 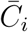 and 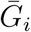 at *t* = 24 h. We see that 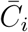 is accurately approximated by 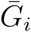 when *D*_*c*_ = 10 and 10^5^ *µ*m^2^/h, whereas the comparison is poor when *D*_*c*_ = 10^6^ *µ*m^2^/h. As a result we have relatively good agreement between the full and reduced averaged density profiles in Figure 6(a)–(e) since 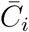 is reasonably well approximated by 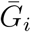. However, results in Figure 6(f) shows that even with a relatively good match between 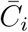 and 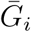, the match between the averaged density profiles of the full and reduced discrete models can still be poor when the strength of chemotaxis is sufficiently large, here *α* = 1000. Results in Figure 6(h)–(i) correspond to cases where 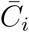 and 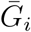 do not match well and in all these cases we see that the average density profiles in the reduced discrete model do not provide a good approximation of the averaged density profiles in the full discrete model.

Overall, comparing the average density profiles in Figure 6 confirms that the reduced discrete model can be used to approximate the full discrete model for certain parameter choices. In general we see that the quality of match between the two models tends to decreases with *α* and the performance of the reduced model is also sensitive to other parameters such as *D*_*c*_. To provide further insight into how the performance of the reduced discrete model depends upon the choice of parameters we compute *E*_1_ and *E*_2_ at *t* = 24 h over a range of *µ, λ, α* and *D*_*c*_. For each choice of *α* and *D*_*c*_, we construct two-dimensional heat maps showing *E*_1_ and *E*_2_ as a function of *µ* and *λ*. The heat maps, shown in Figure 7, indicate that the reduced model provides a reasonably good approximation of the full model provided we have a sufficiently small *α* and *D*_*c*_. Comparing the magnitude of *E*_1_ and *E*_2_ as a function of *λ* and *µ* indicates that the accuracy of the reduced discrete model is less sensitive to variation in *λ* and *µ* than it is to variations in *α* and *D*_*c*_. Similar results (not shown) also indicate that *E*_1_ and *E*_2_ are relatively insensitive to the choice of *κ* for the choice of initial condition to mimic the co–culture experiments in Figure 2. Therefore, we have chosen to focus our examination of the performance of the reduced discrete model to *α, D*_*c*_, *λ* and *µ*.

**Fig. 7.**
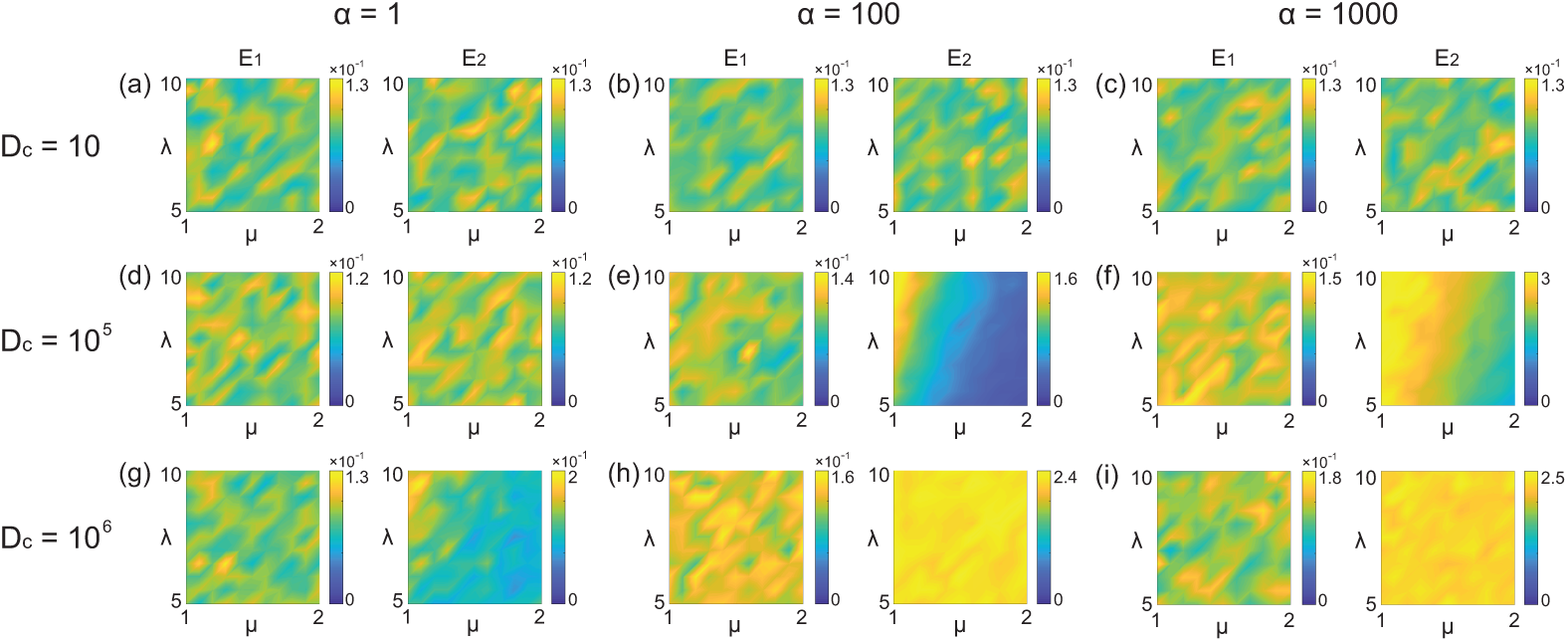
Heat maps showing *E*_1_(24; *D*_*c*_, *α, λ, κ, µ*) and *E*_2_(24; *D*_*c*_, *α, λ, κ, µ*) for various choices of *D*_*c*_, *α, λ* and *µ* with *κ* held constant at *κ* = 1. In all simulations we have Δ = 20 *µ*m and *τ* = 0.01 h.

## 6 Conclusion and Outlook

Typical mathematical models of cell migration stimulated by signalling molecules involve some kind of reaction–diffusion equation to explicitly describe the spatial and temporal distribution of the signalling molecules. However, such information is rarely available from experimental observations since signalling molecules are challenging to record and image. Motivated by a suite of new co– culture cell migration assays, we develop new mathematical modelling tools to describe the cell migration regulated by signalling molecules in an attempt to avoid the need for working directly with a description of the spatial and temporal distribution of signalling molecules. We first develop a full discrete model that describes the migration and interactions of two subpopulations of cells, in which the movement of one subpopulation is regulated by the presence of signalling molecules secreted by cells in the other subpopulation. In this model, the spatial and temporal distribution of the signalling molecules is governed by a discrete conservation statement that is related to a reaction–diffusion equation. To make this description consistent with experimental observations, we simplify the full discrete model by invoking a quasi–steady state assumption in the reaction–diffusion equation governing the spatial and temporal distribution of the signalling molecules. With this simplification, we obtain a reduced discrete model which implicitly describes a similar interaction between the two cell populations without needing to solve the underlying conservation statement. To provide additional mathematical insight into these two models we obtain continuum limit descriptions of both models, leading to new PDE models.

In the full discrete model we suppose that the migration rates of agents from Subpopulation 1 and Subpopulation 2 are given by functions *f*_1_(*C*_*i*_) and *f*_2_(*C*_*i*_), respectively, where *C*_*i*_ is the density of signalling molecules at site *i*. Similarly, in the reduced discrete model the migration rates of agents from Subpopulation 1 and Subpopulation 2 are given by *f*_1_(*G*_*i*_) and *f*_2_(*G*_*i*_), respectively, where *G*_*i*_ is the approximate density of signalling molecules at site *i*. Choosing particular functional forms for *f*_1_ and *f*_2_ allows us to specify whether cell migration is stimulated or inhibited by the signalling molecule. We choose forms of *f*_1_ and *f*_2_ that are relevant to the hepatocype–MSC co–culture experiments in Figure 2, and we compare the performance of the full discrete model and reduced discrete model for a typical experimental geometry, timescale, and parameter choices, and we focus on comparing the full and reduced models for different strengths of the chemokinesis effect. This comparison indicates particular situations where the reduced discrete model could be used in place of the full discrete model. In general we find that the reduced discrete model performs particularly well when the strength of chemokinesis is sufficiently small, whereas for sufficiently strong chemokinesis the comparisons indicate that the reduced model is not always a good approximation. Without making such comparisons, it is not obvious when it would be reasonable to use the reduced model.

There are several features of this study that could warrant further investigation: (i) For simplicity, we focus on developing one–dimensional models to describe cell migration regulated by signalling molecules, and these one– dimensional models can be extended to two–dimensional geometries where necessary; (ii) In all comparisons we assume *C*_*i*_ = 0 at *t* = 0 in the full model description. This assumption is reasonable given that typical experiments do not provide any information about the spatial and temporal distribution of signalling molecules. If, instead, the initial distribution for *C*_*i*_ was known or measurable, all comparisons in this work could be repeated making use of that information; (iii) In this work we choose particular forms of *f*_1_ and *f*_2_ that are relevant to the hepatocyte–MSC co–culture experiments in Figure 2. Other choices of *f*_1_ and *f*_2_ could be made for different co–culture systems as relevant; and (iv) In the current modelling framework we assume that cells sense signalling molecules locally, at the same site *i*. However, in the cell biology literature there have been different hypotheses put forward about non-local sensing over different spatial ranges (Hopkins and Camley 2019). Such non-local sensing could be introduced into our modelling framework by making appropriate adjustments to the discrete models and then examining how these changes manifest in the continuum limit description.

## Acknowledgments

This work is supported by the Australian Research Council (DP170100474) and the National Health and Medical Research Council (APP1126091, APP1141121). WJ is supported by a QUT Vice-Chancellor’s Research Fellowship.

## References

[1] Alarcón T, Byrne HM, Maini PK (2003) A cellular automaton model for tumour growth in inhomogeneous environment. Journal of Theoretical Biology 225:257–274.

[2] Balding D, McElwain DLS (1985) A mathematical model of tumour–induced capillary growth. Journal of Theoretical Biology 114:53–73.

[3] Behar TN, Li YX, Tran HT, Ma W, Dunlap V, Scott C, Barker JL (1996) GABA stimulates chemotaxis and chemokinesis of embryonic cortical neurons via calcium–dependent mechanisms. Journal of Neuroscience 16:1808–1818.

[4] Breward CJW, Byrne HM, Lewis CE (2002) The role of cell-cell interactions in a two-phase model for avascular tumour growth. Journal of Mathematical Biology. 45: 125–152.

[5] Brumley DR, Carrara F, Hein AM, Yawata Y, Levin SA, Stocker R (2019) Bacteria push the limits of chemotactic precision to navigate dynamic chemical gradients. Proceedings of the National Academy of Sciences 116:10792–10797.

[6] Byrne HM, Cave G, McElwain DLS (1998) The effect of chemotaxis and chemokinesis on leukocyte locomotion: A new interpretation of experimental results. Mathematical Medicine and Biology: A Journal of the IMA 15:235–256.

[7] Byrne HM, Owen MR (2004) A new interpretation of the Keller-Segel model based on multiphase modelling. Journal of Mathematical Biology 49:604–626.

[8] Cai AQ, Landman KA, Hughes BD (2006) Modelling directional guidance and motility regulation in cell migration. Bulletin of Mathematical Biology 68:25–52.

[9] Chen Y, Corriden R, Inoue Y, Yip L, Hashiguchi N, Zinkernagel A, Nizet V, Insel PA, Junger WG (2006) ATP release guides neutrophil chemotaxis via P2Y2 and A3 receptors. Science 314:1792–1795.

[10] Chung S, Sudo R, Mack PJ, Wan CR, Vickerman V, Kamm RD (2009) Cell migration into scaffolds under co-culture conditions in a microfluidic platform. Lab on a Chip 9:269–275.

[11] Das AM, Eggermont AM, Ten Hagen TL (2015) A ring barrier-based migration assay to assess cell migration *in vitro*. Nature Protocols 10:904–915.

[12] Flegg JA, Menon SN, Maini PK, McElwain DLS (2015) On the mathematical modeling of wound healing angiogenesis in skin as a reaction–transport process. Frontiers in Physiology 6:262.

[13] Frimberger D, Morales N, Gearhart JD, Gearhart JP, Lakshmanan Y (2006) Human embryoid body–derived stem cells in tissue engineering–enhanced migration in co-culture with bladder smooth muscle and urothelium. Urology 67:1298–1303.

[14] Gruber HE, Somayaji S, Riley F, Hoelscher GL, Norton HJ, Ingram J, Hanley Jr EN (2012) Human adipose–derived mesenchymal stem cells: serial passaging, doubling time and cell senescence. Biotechnic & Histochemistry 87:303–311.

[15] Hillen T, Painter KJ (2009) A user’s guide to PDE models for chemotaxis. Journal of Mathematical Biology 58:183–217.

[16] Hopkins A, Camley BA (2019) Leader cells in collective chemotaxis: Optimality and trade-offs. Physical Review E 100:032417.

[17] Huang S, Brangwynne CP, Parker KK, Ingber DE (2005) Symmetry–breaking in mammalian cell cohort migration during tissue pattern formation: Role of random–walk persistence. Cell Motility and the Cytoskeleton 61:201–213.

[18] Jin W, Penington CJ, McCue SW, Simpson MJ (2016a) Stochastic simulation tools and continuum models for describing two-dimensional collective cell spreading with universal growth functions. Physical Biology 13:056003.

[19] Jin W, Shah ET, Penington CJ, McCue SW, Chopin LK, Simpson MJ (2016b) Reproducibility of scratch assays is affected by the initial degree of confluence: experiments, modelling and model selection. Journal of Theoretical Biology 390:136–145.

[20] Jin W, Penington CJ, McCue SW, Simpson MJ (2017) A computational modelling framework to quantify the effects of passaging cell lines. PLOS One 12:e0181941.

[21] Keller EF, Segel LA (1971) Model for chemotaxis. Journal of Theoretical Biology 30:225–234.

[22] Kirkpatrick B, Nguyen L, Kondrikova G, Herberg S, Hill WD (2010) Stability of human stromal–derived factor–1*α* (CXCL12*α*) after blood sampling. Annals of Clinical & Laboratory Science 40:257–260.

[23] Kucia M, Jankowski K, Reca R, Wysoczynski M, Bandura L, Allendorf DJ, Zhang J, Ratajczak J, Ratajczak MZ (2004) CXCR4–SDF-1 signalling, locomotion, chemotaxis and adhesion. Journal of Molecular Histology 35:233–245.

[24] Liu Z, Klominek J (2004) Chemotaxis and chemokinesis of malignant mesothelioma cells to multiple growth factors. Anticancer Research 24:1625–1630.

[25] Luster AD (1998) Chemokines–chemotactic cytokines that mediate inflammation. New England Journal of Medicine 338:436–445.

[26] Mac Gabhann F, Popel AS (2004) Model of competitive binding of vascular endothelial growth factor and placental growth factor to VEGF receptors on endothelial cells. American Journal of Physiology–Heart and Circulatory Physiology 286:H153–H164.

[27] Mallet DG, de Pillis LG (2006) A cellular automata model of tumor–immune system interactions. Journal of Theoretical Biology 239:334–350.

[28] Murray JD (2002) Mathematical Biology, 3rd edn. Springer, Berlin.

[29] Novo E, Busletta C, di Bonzo LV, Povero D, Paternostro C, Mareschi K, Ferrero I, David E, Bertolani C, Caligiuri A, Cannito S, Tamagno E, Compagnone A, Colombatto S, Marra S, Fagioli F, Pinzani M, Parola M (2011) Intracellular reactive oxygen species are required for directional migration of resident and bone marrow–derived hepatic pro–fibrogenic cells. Journal of Hepatology 54:964–974.

[30] Painter KJ, Maini PK, Othmer HG (2000) Development and applications of a model for cellular response to multiple chemotactic cues. Journal of Mathematical Biology 41:285–314.

[31] Painter KJ, Sherratt JA (2003) Modelling the movement of interacting cell populations. Journal of Theoretical Biology 225:327–339.

[32] Pettet GJ, Byrne HM, McElwain DLS, J Norbury (1996) A model of wound healing angiogenesis in soft rissue. Mathematical Biosciences 136: 35–63.

[33] Pillay S, Byrne HM, Maini PK (2018) The impact of exclusion processes on angiogenesis models. Journal of Mathematical Biology 77:1721–1759.

[34] Richards GR, Millard RM, Leveridge M, Kerby J, Simpson PB (2004) Quantitative assays of chemotaxis and chemokinesis for human neural cells. Assay and Drug Development Technologies 2:465–472.

[35] Rosoff WJ, Urbach JS, Esrick MA, McAllister RG, Richards LJ, Goodhill GJ (2004) A new chemotaxis assay shows the extreme sensitivity of axons to molecular gradients. Nature Neuroscience 7:678–682.

[36] Sherratt JA, Sage EH, Murray JD (1993) Chemical control of eukaryotic cell movement: A new model. Journal of Theoretical Biology 162:23–40.

[37] Sherratt JA (1994) Chemotaxis and chemokinesis in eukaryotic cells: The Keller-Segel equations as an approximation to a detailed model. Bulletin of Mathematical Biology 56:129–146.

[38] Simpson MJ, Landman KA, Newgreen DF (2006) Chemotactic and diffusive migration on a non-uniformly growing domain: Numerical algorithm development and applications. Journal of Computational and Applied Mathematics 192: 282–300.

[39] Simpson MJ, Landman KA, Hughes BD, Fernando AE (2010) A model for mesoscale patterns in motile populations. Physica A: Statistical Mechanics and its Applications. 389: 1412–1424.

[40] Son K, Brumley DR, Stocker R (2015) Live from under the lens: Exploring microbial motility with dynamic imaging and microfluidics. Nature Reviews Microbiology 13:761–775.

[41] Stinner C, Tello JI, Winkler M (2014) Competitive exclusion in a two–species chemotaxis model. Journal of Mathematical Biology 68:1607–1626.

[42] Tokoyoda K, Egawa T, Sugiyama T, Choi BI, Nagasawa T (2004) Cellular niches controlling B lymphocyte behavior within bone marrow during development. Immunity 20:707–718.

[43] Treloar KK, Simpson MJ, McElwain DLS, Baker RE (2014) Are *in vitro* estimates of cell diffusivity and cell proliferation rate sensitive to assay geometry? Journal of Theoretical Biology 356: 71–84.

[44] Veldkamp CT, Ziarek JJ, Su J, Basnet H, Lennertz R, Weiner JJ, Peterson FC, Baker JE, Volkman BF (2009) Monomeric structure of the cardioprotective chemokine SDF-1/CXCL12. Protein Science 18:1359–1369.

[45] Wright LM, Maloney W, Yu X, Kindle L, Collin-Osdoby P, Osdoby P (2005) Stromal cell–derived factor-1 binding to its chemokine receptor CXCR4 on precursor cells promotes the chemotactic recruitment, development and survival of human osteoclasts. Bone 36:840–853.

[46] Yoon D, Kim H, Lee E, Park MH, Chung S, Jeon H, Ahn CH, Lee K (2016) Study on chemotaxis and chemokinesis of bone marrow–derived mesenchymal stem cells in hydrogel-based 3D microfluidic devices. Biomaterials Research 20:25.

