## Supplementary Material for "New mathematical modelling tools for co-culture experiments: when do we need to explicitly account for signalling molecules?"

---

**Contents**


1 Numerical methods 3

1.1 Signalling molecules in the discrete models 3

1.2 Continuum models 4

2 Additional results for chemokinesis over longer timescales 5

3 Discrete models describing chemotaxis 7

3.1 Full discrete model 7

3.2 Reduced discrete model 8

4 Continuum limit descriptions describing chemotaxis 10

4.1 Full continuum model 10

4.2 Reduced continuum model 12

5 Comparison of results for chemotaxis 14

6 Experimental protocols 16

6.1 Cell culture 16

6.2 Ring barrier–based co–culture migration assay 16

6.3 Long–term time–lapse microscopy and analysis of cell migration 16

6.4 Immunofluorescence 17

References 17

### 1 Numerical methods

#### *1.1 Signalling molecules in the discrete models*

The spatial and temporal distribution of signalling molecules is governed by the discrete conservation statement,

$$\frac{\delta C_i}{\tau} = \frac{D_c}{\Delta^2}\left[C_{i-1} - 2C_i + C_{i+1}\right] + \lambda A_i - \kappa C_i B_i - \mu C_i, \tag{S1}$$

where $D_c$ [$\mu$m$^2$/h] is the molecular diffusivity, $\lambda$ [$\mu$M/h] is the secretion rate, $\kappa$ [/h] is the uptake rate and $\mu$ [/h] is the intrinsic decay rate. To update $C_i$ from time $t$ to $t+\tau$, we denote the density of signalling molecules at time $t$ by $C_i(t)$. Then during the next time step of duration $\tau$, we obtain $C_i(t+\tau)$ by numerically solving

$$\begin{aligned}\frac{C_i(t+\tau) - C_i(t)}{\tau} =& \frac{D_c}{\Delta^2}\Big[C_{i-1}(t+\tau) - 2C_i(t+\tau) + C_{i+1}(t+\tau)\Big] \\ &+ \lambda A_i(t) - \kappa B_i(t) C_i(t+\tau) - \mu C_i(t+\tau).\end{aligned} \tag{S2}$$

It is worth noting that in our random sequential updated method, within the time step of duration $\tau$, $C_i(t+\tau)$ is always updated first since both the diffusion and reaction of typical signalling molecules happen much faster than cell migration. Therefore in Equation (S2) we use $A_i(t)$ and $B_i(t)$ to compute $C_i(t+\tau)$. With reflecting boundary conditions at $i=1$ and $i=I$, Equation (S2) can be rearranged to form a system of coupled linear algebraic equations which can be solved using the Thomas algorithm.

#### *1.2 Continuum models*

The full continuum model (Equations (5)-(7)) and the reduced continuum model (Equations (10)-(11)) for describing chemokinesis are solved numerically using a finite difference method. For each governing equation, we discretise the spatial derivatives using a central difference method with a uniform spacing $\delta x$, and the temporal derivatives using a backward Euler method with a uniform time step $\delta t$. Such discretisation leads to a system of coupled non-linear algebraic equations for both the full and reduced models, which are linearised using Picard iteration with absolute convergence tolerance $\epsilon$, and solved sequentially using the Thomas algorithm (Morton and Mayers 2005). The full and reduced continuum models for describing chemotaxis are also solved using the same approach. For all the results of the continuum models presented in the main document as well as the supplementary material, we choose $\delta x = 2$ $\mu$m, $\delta t = 10^{-4}$ h, and $\epsilon = 10^{-5}$ so that our results are grid-independent.

#### 2 Additional results for chemokinesis over longer timescales

We repeat the simulations in Figure 4 and extend the time duration to 48 h so that the two subpopulations have more opportunity to interact. In Figure S1 we show the snapshots from the full and reduced discrete models at $t = 0, 12, 24, 36$, and 48 h (Figure S1(a)-(f)), the full and reduced model comparisons (Figure S1(g)-(i)), and continuum-discrete match for both full and reduced models (Figure S1(j)-(o)). Overall, we observe the similar trends described in Figure 4.

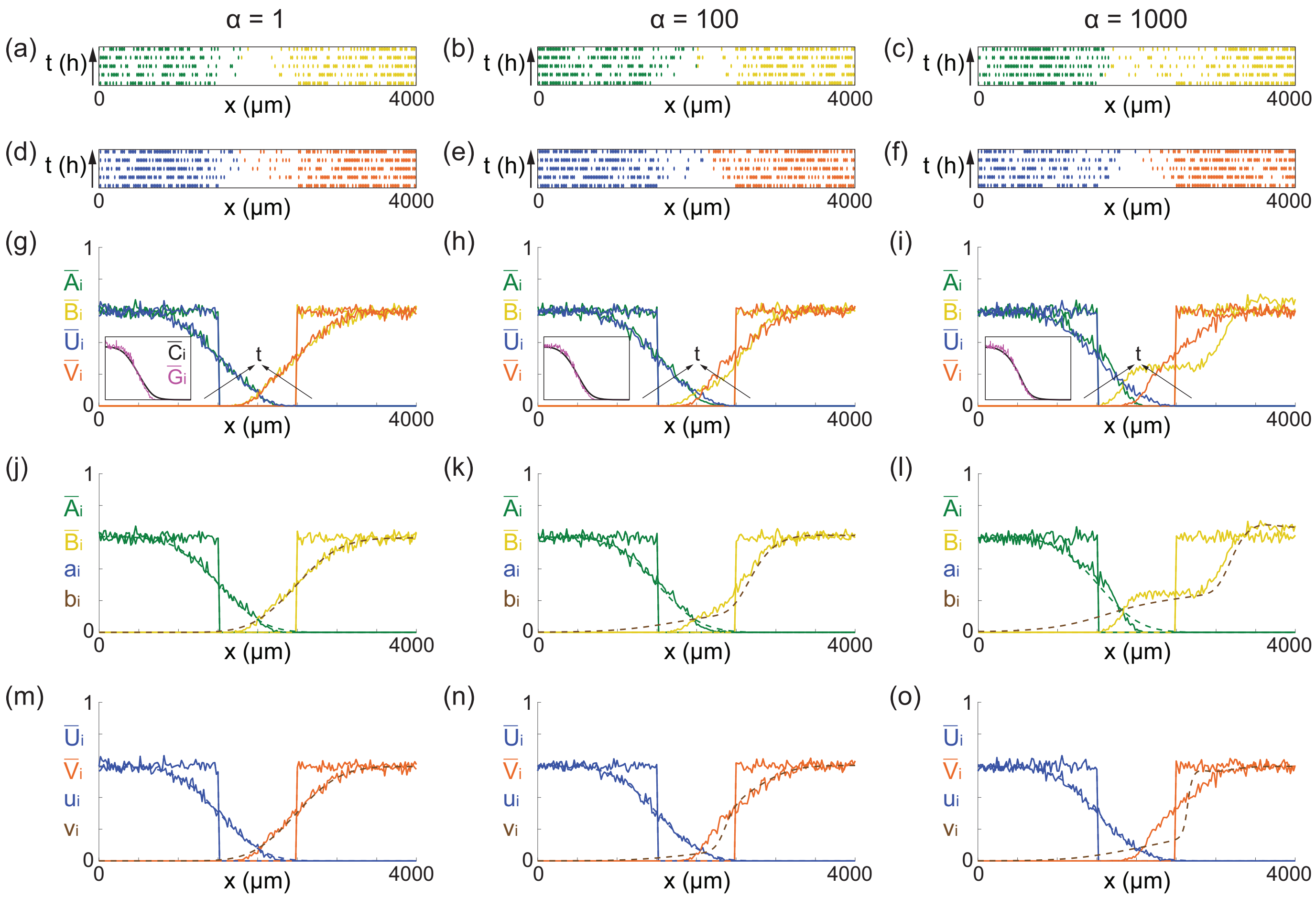


Fig. S1. Comparison of the full and reduced discrete models describing chemokinesis. (a)-(c) Snapshots from the full discrete model at $t = 0, 12, 24, 36,$ and 48 h. The arrow along the vertical direction indicates increasing time. (d)-(f) Snapshots from the reduced discrete model at $t = 0, 12, 24, 36,$ and 48 h. (g)-(i) Density profiles of Subpopulation 1 and Subpopulation 2 from the full and reduced discrete models at $t = 0$ and 48 h. The black arrow indicates increasing time. The inset in each subfigure shows profiles of the $\bar{C}_i$ and $\bar{G}_i$ at $t = 48$ h. (j)-(l) Continuum–discrete comparisons of the full models at $t = 0$ and 48 h. (m)-(o) Continuum–discrete comparisons of the reduced models at $t = 0$ and 48 h. The solid line indicates results from the discrete models. The dashed line indicates results from the continuum models. All the simulation data are obtained by averaging over 500 statistically identically prepared realisations. $D_a = D_b(0) = D_u = D_v(0) = 2000$ $\mu$m$^2$/h, $D_c = 10^5$ $\mu$m$^2$/h, $\lambda = 1$ $\mu$M/h, and $\kappa = \mu = 1$ /h, $\Delta = 20$ $\mu$m and $\tau = 0.01$ h for all the simulations.

#### 3 Discrete models describing chemotaxis

In the main document we present the details of models describing chemokinesis. However, the modelling framework developed in this study can be used to examine either chemokinesis, chemotaxis, or a combination of chemokinesis and chemotaxis. In this section we present the details by focusing on modelling chemotaxis. We adopt the same notations used in the main document for consistency.

##### *3.1 Full discrete model*

The general framework of the full model that describes chemotaxis remains the same as that described in Section 3 in the main document. The full model describing chemotaxis is also implemented using a similar random sequential update method. Within a particular time step of duration $\tau$, after $C_i$ is updated using Equation (S1), $N_1 + N_2$ agents are randomly selected, one at a time, with replacement. Agents from Subpopulation 1 and Subpopulation 2 attempt to undergo a nearest neighbour movement with constant probability $P_{m1}$ and $P_{m2}$, respectively. If an agent from Subpopulation 1 at site $i$ is selected and attempts to move, the agent attempts to move to site $i+1$ with a probability $\Big(1+p_1(\nabla C_i)\Big)/2 \in [0,1]$, where we define $\nabla C_i = C_{i+1} - C_{i-1}$, which is the local difference of the density of signalling molecules. Similarly, if an agent from Subpopulation 2 at site $i$ is selected and attempts to move, the agent attempts to move to site $i+1$ with a probability $\Big(1+p_2(\nabla C_i)\Big)/2 \in [0,1]$. Here both $p_1(\nabla C_i) \in [-1,1]$ and $p_2(\nabla C_i) \in [-1,1]$ are functions depending on the local difference of the density of signalling molecules. We note that at $i=1$ and $i=I$, we have $\nabla C_1 = C_2 - C_1$ and $\nabla C_I = C_I - C_{I-1}$, respectively, due to the reflecting boundary conditions. Potential motility events are aborted if the target site is occupied. Reflecting boundary conditions are applied in all cases we consider.

The functional forms of $p_1(\nabla C_i)$ and $p_2(\nabla C_i)$ determine how the agents respond to the local difference of signalling molecules. For example, if $p_1$ is increasing, agents from Subpopulation 1 are directed up the gradient of the signalling molecule. This process is called chemoattraction. In contrast, if $p_1$ is decreasing, agents from Subpopulaiton 1 are directed down the gradient of the signalling molecule, which is called chemorepulsion. Therefore, $p_1(\nabla C_i)$ and $p_2(\nabla C_i)$ control the attempted direction of migration but do not affect the rate of migration. In this work, we assume that agents from both subpopulations undergo unbiased migration when $\nabla C_i = 0$ and that the signalling molecules have no impact upon the migration of Subpopulation 1 so we set $p_1(\nabla C_i) = 0$. In contrast, we choose $p_2(\nabla C_i)$ to be a smooth increasing function, given by

$$p_2(\nabla C_i) = \frac{1 - \mathrm{e}^{-\beta \nabla C_i}}{1 + \mathrm{e}^{-\beta \nabla C_i}}, \tag{S3}$$

where $\beta \geq 0$ specifies the strength of the chemotactic response. Our choice of $p_1(\nabla C_i)$ and $p_2(\nabla C_i)$ is such that $p_1(\nabla C_i) = p_2(0) = 0$ so that in the absence of the chemical gradient agents from both subpopulations undergo unbiased random migration.

##### *3.2 Reduced discrete model*

We now formulate a reduced discrete model that retains key elements of the full discrete model but avoids the need to explicitly solve for the spatial and temporal distribution of the signalling molecules. Similar to the reduced model describing chemokinesis, we simplify the model by assuming we have localised quasi-steady conditions. Therefore, in our reduced model, $G_i = (\lambda \hat{U}_i)/(\mu + \kappa \hat{V}_i)$ is the approximate density of the signalling molecule density at site $i$. Using this approximation in our discrete modelling framework allows us to implicitly simulate the role of the signalling molecules without needing to solve the underlying reaction–diffusion equation. We further define $\nabla G_i = G_{i+1} - G_{i-1}$ to approximate $\nabla C_i$.

The reduced discrete model is implemented using a similar random sequential update method. The only difference is that in the reduced discrete model the two functions $p_1(\nabla G_i)$ and $p_2(\nabla G_i)$ depend on $\nabla G_i$ instead of $\nabla C_i$, and there is no need to solve the evolution equation for $C_i$.

### 4 Continuum limit descriptions describing chemotaxis

#### 4.1 *Full continuum model*

In this section we show the derivation of the continuum limit description of the full discrete model describing chemotaxis. Following the same approach described in the main document, we invoke a mean-field assumption and accounting for all possible events that alter the occupancy of site $i$ over a time step of duration $\tau$, we obtain

$$\delta\bar{A}_i = \overbrace{\frac{P_{m1}}{2}\left(1-\bar{S}_i\right)\left[\left(1+p_1(\nabla C_{i-1})\right)\bar{A}_{i-1} + \left(1-p_1(\nabla C_{i+1})\right)\bar{A}_{i+1}\right]}^{\text{increase in occupancy due to migration into site } i}$$
$$- \overbrace{\frac{P_{m1}}{2}\bar{A}_i\left[\left(1-p_1(\nabla C_i)\right)\left(1-\bar{S}_{i-1}\right) + \left(1+p_1(\nabla C_i)\right)\left(1-\bar{S}_{i+1}\right)\right]}^{\text{decrease in occupancy due to migration out of site } i}, \tag{S4}$$

$$\delta\bar{B}_i = \overbrace{\frac{P_{m2}}{2}\left(1-\bar{S}_i\right)\left[\left(1+p_2(\nabla C_{i-1})\right)\bar{B}_{i-1} + \left(1-p_2(\nabla C_{i+1})\right)\bar{B}_{i+1}\right]}^{\text{increase in occupancy due to migration into site } i}$$
$$- \overbrace{\frac{P_{m2}}{2}\bar{B}_i\left[\left(1-p_2(\nabla C_i)\right)\left(1-\bar{S}_{i-1}\right) + \left(1+p_2(\nabla C_i)\right)\left(1-\bar{S}_{i+1}\right)\right]}^{\text{decrease in occupancy due to migration out of site } i}, \tag{S5}$$

where $\delta\bar{A}_i$ and $\delta\bar{B}_i$ are the change in occupancy at site $i$ of Subpopulation 1 and 2, respectively, and $\bar{S}_i = \bar{A}_i + \bar{B}_i$ is the total average occupancy at site $i$. To convert these discrete conservation statements into continuous expression we identify the discrete variables with continuous variables, $\bar{A}_i(t) = a(x,t)$, $\bar{B}_i(t) = b(x,t)$ and $\bar{C}_i(t) = c(x,t)$ . Expanding each term in Equations (S4)–(S5) about site $i$ using a Taylor series and neglecting terms of $\mathcal{O}(\Delta^3)$, we divide both sides of the resulting expressions by $\tau$ and take the limit $\Delta \to 0$

and $\tau \to 0$ jointly, with $\Delta^2/\tau$ held constant, to give

$$\frac{\partial a}{\partial t} = D_a \frac{\partial}{\partial x}\left[(1-s)\frac{\partial a}{\partial x} + a\frac{\partial s}{\partial x}\right] - \chi_a \frac{\partial}{\partial x}\left[(1-s)\,a\frac{\partial c}{\partial x}\right] - \omega_a \frac{\partial}{\partial x}\left[(1-s)\,a\right], \tag{S6}$$

$$\frac{\partial b}{\partial t} = D_b \frac{\partial}{\partial x}\left[(1-s)\frac{\partial b}{\partial x} + b\frac{\partial s}{\partial x}\right] - \chi_b \frac{\partial}{\partial x}\left[(1-s)\,b\frac{\partial c}{\partial x}\right] - \omega_b \frac{\partial}{\partial x}\left[(1-s)\,b\right], \tag{S7}$$

$$\frac{\partial c}{\partial t} = D_c \frac{\partial^2 c}{\partial x^2} + \lambda a - \kappa c b - \mu c, \tag{S8}$$

where $D_a = \Delta^2 P_{m1}/(2\tau)$ and $D_b = \Delta^2 P_{m2}/(2\tau)$ are the diffusion coefficients for Subpopulation 1 and Subpopulation 2, respectively; $\chi_a = 2p_1'(0)P_{m1}\Delta^2/\tau$ and $\chi_b = 2p_2'(0)P_{m2}\Delta^2/\tau$ are the chemotaxis coefficients of Subpopulation 1 and Subpopulation 2, respectively; $\omega_a = p_1(0)P_{m1}\Delta/\tau$ and $\omega_b = p_2(0)P_{m2}\Delta/\tau$ are the drift velocities of Subpopulation 1 and Subpopulation 2, respectively; and $s(x,t) = a(x,t)+b(x,t)$. We note that both $p_1(0) = \mathcal{O}(\Delta)$ and $p_2(0) = \mathcal{O}(\Delta)$ are required in order to obtain a well–defined continuum limit (Simpson et al. 2010).

It is worth noting that in the full discrete model both $p_1(\nabla C_i)$ and $p_2(\nabla C_i)$ are functions of the local difference of signalling molecule density, while in the continuum limit descriptions, $p_1(\partial C/\partial x)$ and $p_2(\partial C/\partial x)$ are functions of the gradient of signalling molecules. This is consistent with the concept of chemotaxis, such that cells sense the gradient of signalling molecules instead of their local density.

Although there are three grouped terms at the right side of both Equations (S6)-(S7), the second term in both equations vanish when $p_1(\partial C/\partial x)$ and $p_2(\partial C/\partial x)$ are constant since $p_1'(0) = p_2'(0) = 0$. The resulting model relaxes to a simpler model describing biased cell migration with constant drift developed previously (Simpson et al. 2009). In contrast, when both $p_1(\partial C/\partial x)$ and $p_2(\partial C/\partial x)$ are not constant, the third term on the right of Equations (S6)-

(S7) vanishes since $p_1(0) = p_2(0) = 0$. Under this circumstance, the resulting model has a similar form of the generalised Keller–Segel model for chemotaxis (Hillen and Painter 2009).

##### *4.2 Reduced continuum model*

The continuum limit description of the reduced discrete model can be obtained using a very similar approach. The approximate conservation statements for the two subpopulations can be written as,

$$\delta\bar{U}_i = \overbrace{\frac{P_{m1}}{2}\left(1-\bar{S}_i\right)\left[\left(1+p_1(\nabla G_{i-1})\right)\bar{U}_{i-1} + \left(1-p_1(\nabla G_{i+1})\right)\bar{U}_{i+1}\right]}^{\text{increase in occupancy due to migration into site } i}$$
$$- \overbrace{\frac{P_{m1}}{2}\bar{U}_i\left[\left(1-p_1(\nabla G_i)\right)\left(1-\bar{S}_{i-1}\right) + \left(1+p_1(\nabla G_i)\right)\left(1-\bar{S}_{i+1}\right)\right]}^{\text{decrease in occupancy due to migration out of site } i}, \tag{S9}$$

$$\delta\bar{V}_i = \overbrace{\frac{P_{m2}}{2}\left(1-\bar{S}_i\right)\left[\left(1+p_2(\nabla G_{i-1})\right)\bar{V}_{i-1} + \left(1-p_2(\nabla G_{i+1})\right)\bar{V}_{i+1}\right]}^{\text{increase in occupancy due to migration into site } i}$$
$$- \overbrace{\frac{P_{m2}}{2}\bar{V}_i\left[\left(1-p_2(\nabla G_i)\right)\left(1-\bar{S}_{i-1}\right) + \left(1+p_2(\nabla G_i)\right)\left(1-\bar{S}_{i+1}\right)\right]}^{\text{decrease in occupancy due to migration out of site } i}, \tag{S10}$$

where all terms have a similar interpretation to those in Equations (S4)–(S5). We proceed to the continuum limit in the same way, arriving at

$$\frac{\partial u}{\partial t} = D_u\frac{\partial}{\partial x}\left[\left(1-s\right)\frac{\partial u}{\partial x} + u\frac{\partial u}{\partial x}\right] - \chi_u\frac{\partial}{\partial x}\left[\left(1-s\right)u\frac{\partial g}{\partial x}\right] - \omega_u\frac{\partial}{\partial x}\left[\left(1-s\right)u\right], \tag{S11}$$

$$\frac{\partial v}{\partial t} = D_v\frac{\partial}{\partial x}\left[\left(1-s\right)\frac{\partial v}{\partial x} + v\frac{\partial v}{\partial x}\right] - \chi_v\frac{\partial}{\partial x}\left[\left(1-s\right)v\frac{\partial g}{\partial x}\right] - \omega_v\frac{\partial}{\partial x}\left[\left(1-s\right)v\right]. \tag{S12}$$

where $u(x,t)$ and $v(x,t)$ are the densities of Subpopulation 1 and Subpopulation 2, respectively. Here, $D_u = \Delta^2 P_{m1}/(2\tau)$ and $D_v = \Delta^2 P_{m2}/(2\tau)$ are the diffusion coefficients of Subpopulation 1 and Subpopulation 2, respectively; $\chi_u = 2p_1'(0)P_{m1}\Delta^2/\tau$ and $\chi_v = 2p_2'(0)P_{m2}\Delta^2/\tau$ are the chemotaxis coefficients of Subpopulation 1 and Subpopulation 2, respectively; $\omega_u = p_1(0)P_{m1}\Delta/\tau$ and $\omega_v = p_2(0)P_{m2}\Delta/\tau$ are the drift velocities of Subpopulation 1 and Subpopulation 2, respectively. Again, both $p_1(\nabla G_i)$ and $p_2(\nabla G_i)$ in the discrete model are associated with $p_1(\partial G/\partial x)$ and $p_2(\partial G/\partial x)$ in the continuum limit descriptions. We require $\Delta \to 0$ and $\tau \to 0$ jointly, with $\Delta^2/\tau$ held constant, and both $p_1(0) = \mathcal{O}(\Delta)$ and $p_2(0) = \mathcal{O}(\Delta)$ for the continuum limit to be well–defined (Simpson et al. 2010).

#### 5 Comparison of results for chemotaxis

We solve the full and reduced discrete models describing chemotaxis, as well as the associated continuum models, using the same initial and boundary conditions proposed in the main document. In addition, we set $D_c = 10^5$ $\mu$m$^2$/h, $\lambda = 1$ $\mu$M/h, and $\mu = \kappa = 1$ /h.

Results in Figure S2(a),(d) show snapshots of the time evolution of agent positions in the full and reduced discrete models, respectively. In these preliminary simulations we specify a weak chemotactic effect, $\beta = 1$. Comparing the distribution of agents in different rows of the subfigures shows that the two subpopulations migrate into the initially–vacant space over time. We estimate the expected behaviour of the simulations by averaging the occupancy of each lattice site using 500 identically–prepared realisations of the stochastic models and show the averaged density profiles in Figure S2(g) where we see that the averaged density profiles from the reduced discrete model compares very well when $\beta$ is small. However, comparing the averaged density profiles in Figure S2(i) illustrates that the reduced discrete model does not approximate the full discrete model very well when the $\beta$ is very large.

Results in Figure S2(j)–(l) compare averaged density profiles from the full model with corresponding solutions of Equations (S6)–(S8) for $\beta = 1, 100$ and 1000, respectively. These results show that the new PDE models provide a good approximation of the averaged behaviour of the full discrete model when $\beta = 1$ and $\beta = 100$, but that the solution of the continuum limit PDE does not provide an accurate approximation of the averaged data from the full discrete model when chemokinesis is sufficiently strong, $\beta = 1000$. Similarly, results in Figure S2(m)–(o) compare averaged density profiles from the reduced model with corresponding solutions of Equations (S11)–(S12) for $\beta = 1, 100$ and 1000, respectively. Again, the same trend is observed from these results.

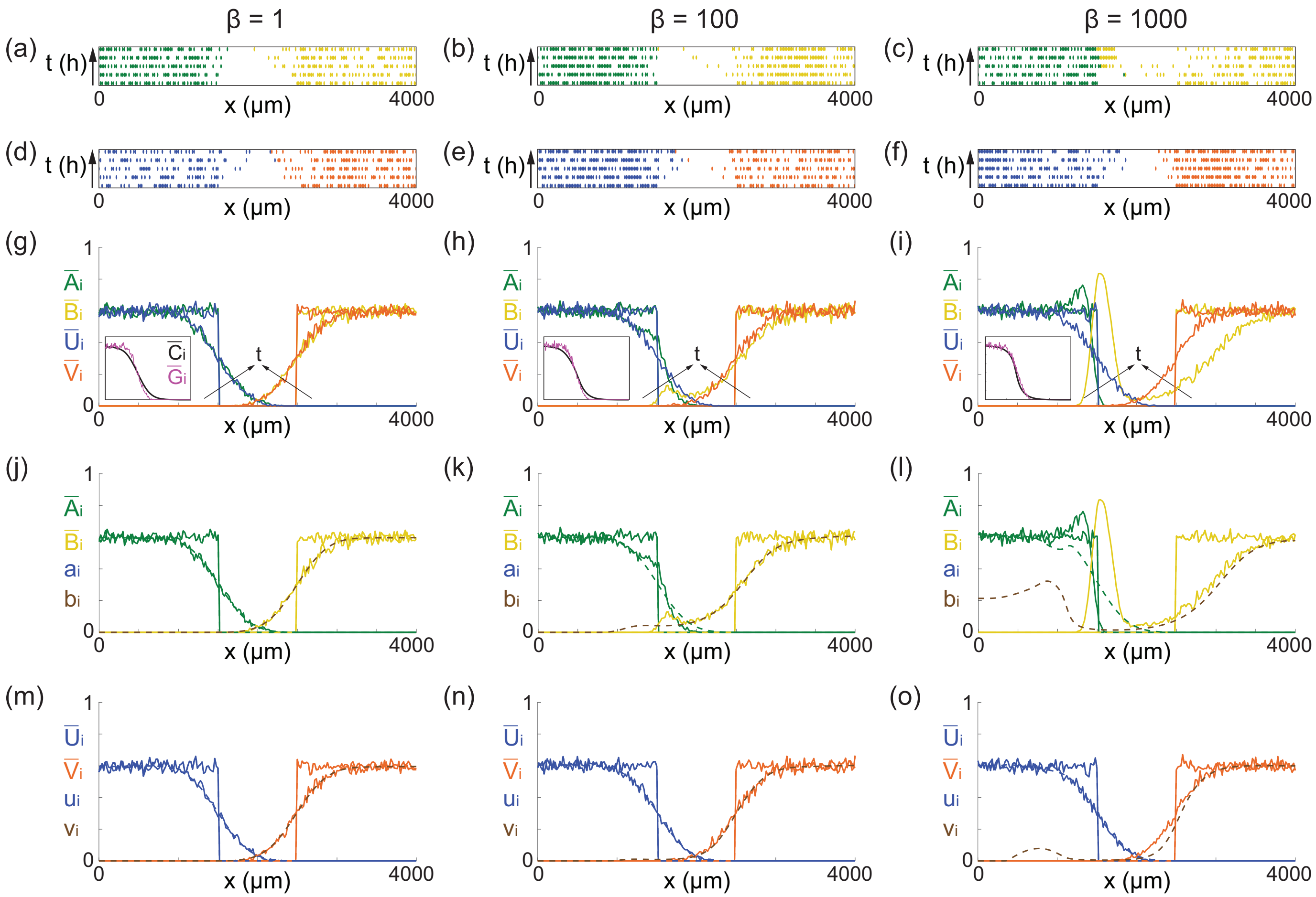


Fig. S2. Comparison of the full and reduced discrete models describing chemotaxis. (a)-(c) Snapshots from the full discrete model at $t = 0, 6, 12, 18,$ and 24 h. The arrow along the vertical direction indicates increasing time. (d)-(f) Snapshots from the reduced discrete model at $t = 0, 6, 12, 18,$ and 24 h. (g)-(i) Density profiles of Subpopulation 1 and Subpopulation 2 from the full and reduced discrete models at $t = 0$ and 24 h. The black arrow indicates increasing time. The inset in each subfigure shows profiles of the $\bar{C}_i$ and $\bar{G}_i$ at $t = 24$ h. (j)-(l) Continuum–discrete comparisons of the full models at $t = 0$ and 24 h. (m)-(o) Continuum–discrete comparisons of the reduced models at $t = 0$ and 24 h. The solid line indicates results from the discrete models. The dashed line indicates results from the continuum models. All the simulation data are obtained by averaging over 500 statistically identically prepared realisations. $D_a = D_b(0) = D_u = D_v(0) = 2000\ \mu\text{m}^2/\text{h}$, $D_c = 10^5\ \mu\text{m}^2/\text{h}$, $\lambda = 1\ \mu\text{M/h}$, and $\kappa = \mu = 1$ /h, $\Delta = 20\ \mu\text{m}$ and $\tau = 0.01$ h for all the simulations.

#### 6 Experimental protocols

##### *6.1 Cell culture*

Cultures of human bone marrow mesenchymal stem/stromal cells (MSCs) are established as previously described (Jin et al. 2018). The human hepatocyte clonal PH5CH8 are kindly provided by Dr Jason C. Steel (School of Health, Medical and Applied Sciences, Central Queensland University, Australia) and cultured in monolayer using high glucose DMEM (ThermoFischer) supplemented with 10% fetal bovine serum (FBS; Thermo Fisher, Waltham, MA, USA) in 95% air and 5% $CO_2$ incubator.

##### *6.2 Ring barrier–based co–culture migration assay*

Ring barrier–based co–culture migration assays are performed using the ring-barrier migration assay previously described (Das et al. 2015), with minor modifications. Briefly, a removable circular sterile migration barrier is inserted into the chamber, which averts cell growth in the center of the coated coverslip. $8 \times 10^5$ MSCs are seeded outside this barrier and incubated at 37 °C for 24 h. Post 24 h, $6 \times 10^3$ PH5CH8 cells are stained with CMPTX (Thermo Fisher Scientific, USA) and seeded inside this ring–barrier and incubated at 37 °C for another 24 h. the migration barrier is removed and the cells are washed twice with PBS, followed by incubation with high glucose DMEM supplemented with 10% FBS for migration assays.

##### *6.3 Long–term time–lapse microscopy and analysis of cell migration*

Time–lapse imaging is conducted on Olympus IX–81 inverted fluorescence microscope (Olympus, Mount Waverley, VIC, Australia). The cell migration is monitored for 24 h. The cells in the incubation chamber are maintained at 37 °C in a constantly humidified atmosphere, with controlled and heated $CO_2$ flow. Images of migrating cells are captured every 30 min, for the duration of 24 h, using a 10X objective (Carl Zeiss). Time–lapse movies are used to quantify parameters of cell migration as previously described (Das et al. 2015).

##### *6.4 Immunofluorescence*

Cells are fixed with 4% Paraformaldehyde for 15 min at room temperature and permeabilised with 0.5% Triton X–100 for 15 min, 3% BSA–PBS for 30 min at room temperature. Primary CXCR4 antibody (Abcam, ab124824, 1/100 dilution) is diluted in 1% BSA–PBS, and incubated overnight at 4 °C. Secondary antibody (Abcam, ab150077) is diluted in 1% BSA–PBS and incubated for 1 h at room temperature. Cell nuclei are stained with DAPI and cells are mounted in Fluoromount medium (Sigma F4680).

#### References

[1] Das AM, Eggermont AM, Ten Hagen TL (2015) A ring barrier–based migration assay to assess cell migration *in vitro*. Nature Protocols 10:904-915.

[2] Hillen T, Painter KJ (2009) A user's guide to PDE models for chemotaxis. Journal of Mathematical Biology 58:183-217.

[3] Jin W, Liang X, Brooks A, Futrega K, Liu X, Doran MR, Simpson MJ, Roberts MS, Wang H (2018) Modelling of the SDF-1/CXCR4 regulated *in vivo* homing of therapeutic mesenchymal stem/stromal cells in mice. PeerJ 6:e6072.

[4] Morton KW, Mayers DF (2005) Numerical Solution of Partial Differential Equations. Cambridge University Press, Cambridge.

[5] Simpson MJ, Landman KA, Hughes BD (2009) Distinguishing between directed and undirected cell motility within an invading cell population. Bulletin of Mathematical Biology 71:781-799.

[6] Simpson MJ, Landman KA, Hughes BD (2010) Cell invasion with proliferation mechanisms motivated by time-lapse data. Physica A: Statistical Mechanics and its Applications 389:3779-3790.
